# MatK impacts Differential Chloroplast Translation of Ribosomal and Photosynthetic genes by limiting spliced tRNA-K(UUU) abundance

**DOI:** 10.1101/2023.10.25.563914

**Authors:** Jose M. Muino, Yujiao Qu, Hannes Ruwe, Sascha Maschmann, Wei Chen, Reimo Zoschke, Uwe Ohler, Kerstin Kaufmann, Christian Schmitz-Linneweber

## Abstract

The protein levels of chloroplast photosynthetic genes and genes related to the chloroplast genetic apparatus vary to adapt to different conditions. However, the underlying mechanisms governing these variations remain unclear. The chloroplast intron Maturase K is encoded within the *trnK* intron and has been suggested to be required for splicing several group IIA introns, including the *trnK* intron. In this study, we employed RNA immunoprecipitation followed by high-throughput sequencing (RIP-Seq) to identify MatK’s preference for binding to group IIA intron domains I and VI within target transcripts. Importantly, these domains are crucial for branch point selection, and we discovered alternative branch points in three MatK target introns, the first observed instances of alternative splicing in chloroplasts. The alternative *trnK* lariat structure showed increased accumulation during heat acclimation. The cognate codon of tRNA-K(UUU) is highly enriched in mRNAs encoding ribosomal proteins and ribosome profiling in a *trnK-matK* over-expressor exhibited elevated levels of the spliced tRNA-K(UUU). Our analysis revealed a significant up-shift in the translation of ribosomal proteins compared to photosynthetic genes. Our findings suggest the existence of a novel regulatory mechanism linked to the abundance of tRNA-K(UUU), enabling the differential expression of functional chloroplast gene groups.

## Introduction

Chloroplast genomes in land plants contain two main functional categories of genes: those that directly encode components of the photosynthetic apparatus (referred to as PS genes) and those that encode components of the genetic apparatus (referred to as GA genes). These two sets of genes have been shown to exhibit differential gene expression during leaf development. In maize, for example, the genes responsible for the ribosome and the RNA polymerase exhibit strong translation in the leaf base where proplastids are located and chloroplast biogenesis begins. Conversely, the translation of photosynthetic genes occurs at a later stage (1). Similar observations have been made in barley and Arabidopsis (2–4). This expression pattern potentially aids in priming the gene expression apparatus in proplastids for the subsequent massive production of photosynthetic proteins.

The mechanistic basis of this differential translation remains unclear, but both transcriptional and post-transcriptional processes are known to play a role (1). It can be expected that the differential expression of these two gene groups is driven, at least in part, by differential transcription. Most GA genes possess promoters for a nuclear-encoded RNA polymerase (NEP), while PS genes are predominantly transcribed by the plastid-encoded RNA polymerase (PEP). Although both RNA polymerases are active in various tissues, NEP-dependent activity is typically strong in young leaf tissue, whereas PEP is active throughout the leaf (5). In addition to transcription, post-transcriptional processes, particularly translation, are believed to contribute significantly to the regulation of chloroplast genes (6).

Chloroplast RNA metabolism is characterized by a variety of RNA processing events, including RNA cleavage, RNA editing, and RNA splicing (7). This complexity is paralleled by the large number of RNA processing factors that have been identified to date. Several families of nuclear-encoded RNA binding proteins have evolved to specifically regulate plant organellar RNA metabolism (7). More than 150 factors have been described as necessary for chloroplast RNA metabolism alone (7). Apart from these nuclear-encoded factors, only one chloroplast-encoded putative RNA processing factor, named Maturase K (MatK), has been identified. MatK is related to bacterial intron maturases and is found in all chloroplast genomes of autotrophically growing land plants, as well as in charophycean green algae (8). It is located within the *trnK* intron and is believed to be involved in splicing its own intron, similar to bacterial maturases. However, in certain species where the *trnK* gene is absent, such as streptophyte algae *Zygnema*, the fern *Adiantum capillus-veneris*, and parasitic land plants (*Epifagus virginiana*, several *Cuscuta* species), *matK* exists as an independent reading frame (9–13). The fact that the loss of *trnK* does not correspond to the loss of *matK* suggests that *matK* serves other functions and potentially targets different introns. Only *Rhizanthella gardneri*, a mycoheterotrophic orchid, and certain members of the *Cuscuta* subgenus of parasitic angiosperms have been observed to lack *matK*. These species have also lost several group IIA introns that may require the activity of MatK (12–14). The available evidence supports the hypothesis that MatK is involved in splicing other introns, not just its own intron. This is further supported by studies on chloroplasts devoid of a translational apparatus, where the *trnK* precursor RNA is not spliced (15) and in addition, an entire subgroup of chloroplast introns, termed group IIA introns, fails to splice as well (16, 17). The only plausible factor that would require functional chloroplast translation for splicing is MatK, leading to the proposition that MatK is responsible for splicing all group IIA introns. Indeed, MatK has been shown to interact with intron RNA both *in vitro* (18) and *in vivo* (19), and recombinant MatK supports splicing in an *in vitro* splicing assay (20).

Group II introns are characterized by six secondary structure elements, named domain DI-DVI, which fold into a globular tertiary structure (21, 22). Bacterial maturase reading frames, including *matK*, are always encoded within DIV. Bacterial maturases establish direct contacts with selected intron sequence elements, and these contacts are essential for the splicing reaction. For example, the bacterial Maturase LtrA interacts with sequence stretches within DI, DII, and DIV. These contacts contribute to the attainment of a splicing-competent intron conformation (23, 24). However, little is known about the preferences of MatK for specific intron domains or how MatK supports intron splicing in living cells.

Functional genetic studies of MatK have been hindered so far by the inability to disrupt the chloroplast *matK* reading frame through various mutagenesis approaches. This suggests that *matK* may be an essential gene (19, 25). A natural mutation in MatK of *Cryptomeria japonica* remains heteroplastomic and leads to infrequent segregation of chlorophyll-deficient sectors, further supporting the notion that MatK is essential for chloroplast development and the survival of whole plant cells (26). Similarly, an attempt to ectopically overexpress MatK from the tobacco chloroplast genome resulted in variegated plants with white sectors and impaired chloroplast development (27).

In this study, we identify the specific interaction sites of MatK with chloroplast introns and demonstrate that MatK is required for the correct branch-point selection of the *trnK* intron. Furthermore, we show that *trnK* production correlates with the translation of GA genes, which are rich in lysine codons. Overexpression of MatK induces increased GA translation, likely mediated by increased *trnK* accumulation, a mechanism that appears to be relevant during leaf development and heat acclimation.

## Results

### Comprehensive Analysis of Binding Sites within Group II A Introns identifies a preference of MatK for domains I and VI

The resolution of prior RIP-Chip analyses investigating the RNA targets of the MatK protein was limited by probe size (19). In order to gain more precise insights into MatK binding sites within its target introns, we employed RNA Immunoprecipitation Sequencing (RIP-Seq). To precipitate MatK, we utilized transplastomic *Nicotiana tabacum* plants expressing MatK with a C-terminal Hemagglutinin (HA) epitope tag (+HA), alongside a non-tagged control line (-HA) as previously described (19). The RNA co-precipitated with MatK was examined via RNA-seq, with the objective to determine the MatK binding sites more precisely. We sequenced total chloroplast RNA from *N. tabacum* as input control for the RIP-seq experiment.

In light of the fact that MatK protein expression peaks in 7-day-old tobacco plants (Zoschke et al., 2010), we isolated chloroplasts from seedlings of this exact age for our analysis. Moreover, given that MatK is a soluble protein located in the chloroplast stroma (Zoschke et al., 2010), we separated the stroma fraction from the membrane fraction to increase the concentration of MatK protein. Consequently, immunoprecipitation (IP) was carried out using the stroma fraction, with the effectiveness of this approach confirmed through immunoblot analysis (Figure 1A). We then extracted and sequenced the RNA co-precipitated with MatK from three independent biological replicates, and determined the relative enrichment compared to IPs from-HA plants. As anticipated, the most pronounced relative enrichment was observed for the already known intron targets of MatK (Figure S1; 19; the intron in trnV is an exception as it does not appear to be enriched here).There were no peaks for RNAs containing other intron types or for RNAs containing no intron at all (Figure 1B), confirming the known specificity of MatK (19).

**Figure 1:**
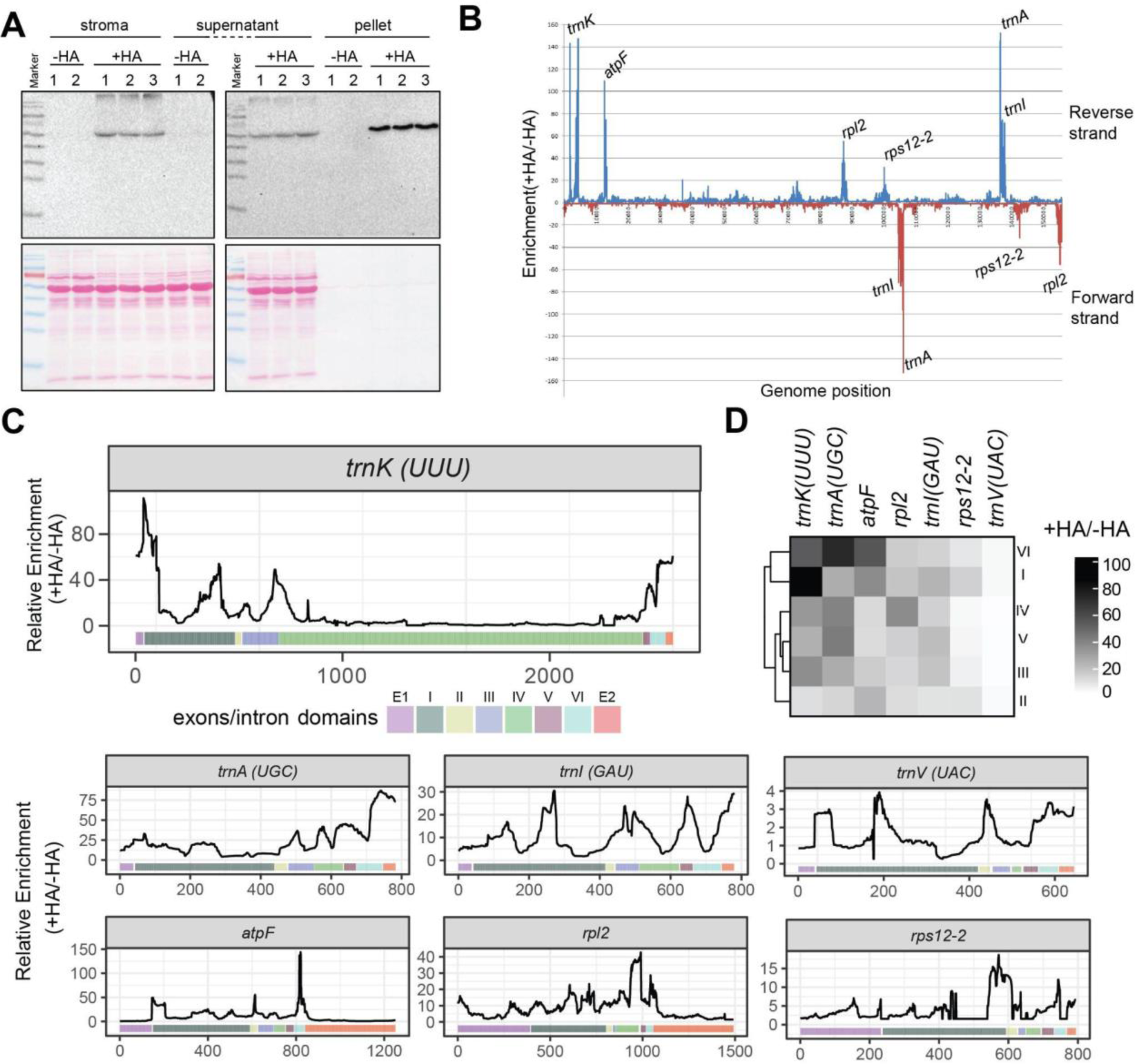
Immunoprecipitation of HA-tagged MatK from transplastomic tobacco lines. A) Chloroplasts were obtained from 7-day-old tobacco seedlings expressing either the HA-tagged MatK (+HA) or the non-tagged control (-HA). Immunoprecipitation of the chloroplast stroma was performed using an HA antibody (Sigma) and magnetic beads (Life Technologies). The protein samples, including the input, IP supernatant, and IP pellet, were separated using SDS-PAGE gel and transferred to a nitrocellulose membrane. HA-specific signals were detected using an HA antibody (upper panel). Ponceau staining was utilized as a loading control (lower panel). The band’s position at about 55 kDa aligns with previous immunological studies on MatK reported by Liere and Link (1995), Barthet and Hilu (2007), and Zoschke et al. (2010). B) Close-up view of the MatK targets identified through RIP-seq. The reads from the +HA and -HA tagged MatK libraries were separately aligned to the chloroplast genome (NC_001879), and the ratio of mapped reads between the +HA and -HA samples was calculated and plotted along the chloroplast genome sequence of the genes shown. The bar below the graph indicates the distribution of exons and domains of group II introns, as previously defined (Michel 1989). The *trnK* plot is enlarged relative to the other introns shown since the intron is far longer because of the *matK* reading frame being positioned within intron domain IV. C) Relative enrichment of each domain of MatK associated RNAs based on RIP-Seq analysis. For each MatK-associated intron, the twenty bases with the highest relative enrichment for each domain were selected and a mean enrichment value (+HA/-HA) was calculated and transformed into a heat map. Based on these peak enrichment values, the domains were clustered using Euclidean distance. D) Comprehensive mapping of MatK targets identified through RIP-seq. RNA samples from the IP pellet were used to generate libraries, which were subsequently sequenced. The resulting reads from each library, corresponding to the +HA and -HA tagged MatK-IP pellets, were mapped to the chloroplast genome (NC_001879). The ratio of mapped reads between the +HA and -HA samples was calculated and plotted onto the chloroplast genome. Peak enrichments are labeled.

The different intron domains fulfill different functions in the splicing process and it is therefore informative to assign binding of MatK to individual domains. Thus, we centered our analysis on the enrichment of RNA across the different intron domains (Figure 1C). Visual inspection of enrichment pointed to domain I and the 3’-terminal domain VI as frequent targets of MatK in almost all introns. For domain I, in most cases peak enrichment is near the immediate 5’-end of the domain as well as in the center of the domain. Additional target sites are found in the other domains (e.g. domain II for *atpF*), but these are not showing higher enrichment compared to binding sites within domain I and VI. Finally, there are binding sites in the second exon of most targets. Since the exons are very short for the four tRNA targets, it is possible that there is simply co-enrichment with domain VI, since the longer second exons of *atpF* and *rpl2* do only show enrichment right adjacent to domain VI within the intron and at much lower enrichment value (Figure 1B).

We next quantified the peak enrichment within each domain – we calculated the mean of the forty bases with the highest enrichment value for each domain and translated this into a heat map (Figure 1C). This focus on top peak height in each domain also avoids the introduction of a length bias, which otherwise leads to an artificial overrepresentation of domain I. We clustered the domains and found that domains I and VI are most similar in their enrichment across all introns and show highest enrichment for most introns. While there appears to be flexibility in target site preferences between introns, our findings indicate a predilection of MatK towards domains I and VI.

### Identification of Alternative Lariat Formation in Chloroplast Introns

Domain VI, a primary target of MatK, plays a crucial role in splicing chemistry. It houses the looped-out adenosine that serves as the branch point during the first splicing phase (28, 29). The selection of the branch point is governed by the structural positioning of Domain VI (DVI) within the group II intron holostructure. This positioning depends on the interactions of exon binding sites in Domain I (DI) that base-pair with exon sequences, thereby aligning the 5’-splice site with the branch point (22). In spliceosomal introns, which have evolved from group II introns, branch point selection relies on the recognition of sequence context by RNA binding proteins and small nuclear ribonucleoproteins (snRNPs). This process can vary, leading to alternative branch points and, therefore, alternative splicing. In chloroplasts, however, alternative splicing has not been reported to date.

We screened standard total RNA-seq datasets from tobacco (30) and Arabidopsis (Zhang et al. Plant Cell 2019) for reads that could help to identify the exact branch point position, thereby determining the possibility of alternative branch points. Such reads would originate immediately downstream of the 5’-splice site and continue across the 2’-5’-phosphodiester linkage at the branch point into the 3’-sequence of the intron (Figure 2A). Our analysis of RNA-seq data from tobacco, Arabidopsis, and maize identified such reads, representing the expected 5’-splice site to known branch point connections for introns in *trnG-UCC*, *petD*, and the two introns in *ycf3* (Figure 2B). Unexpectedly, we also discovered alternative lariats for the introns *trnK*, *trnA*, and *atpF*, all of which are MatK targets (Figure 2C).

**Figure 2:**
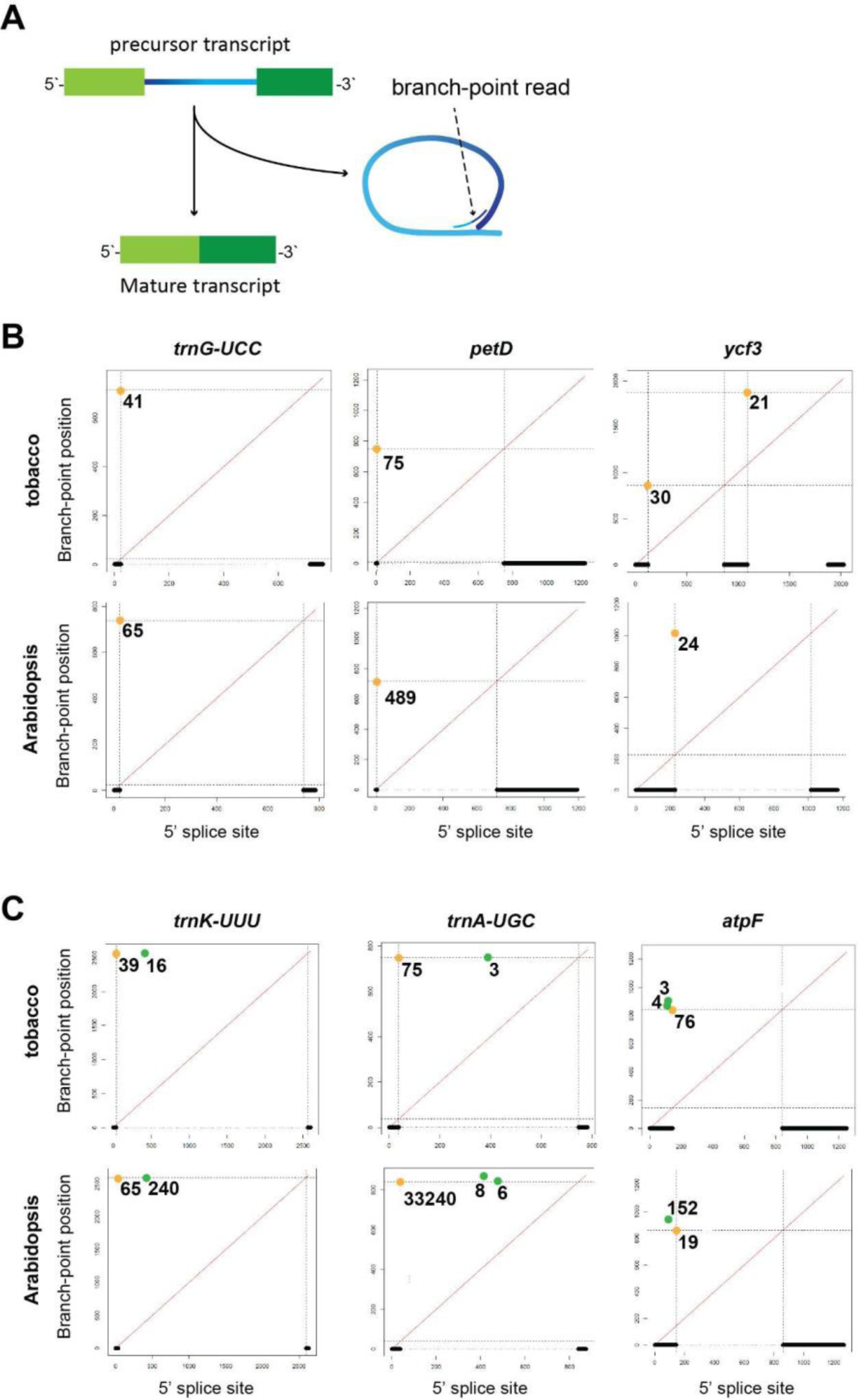
Lariat detection in chloroplast introns in RNA-seq data. A) Graphic representation of reads spanning the 2’-to-5’ bridge at branch-points within lariats. Light green: 5’-exon; dark green: 3’-exon. blue: intron. B) RNA-seq data from tobacco or Arabidopsis were screened for informative reads spanning the 2’-to-5’ bridge at branch-points within lariats. Introns with canonical branch points only: Mapping of branch point reads relative to 5’ splice acceptor sites in tobacco and Arabidopsis. Thick lines at the bottom of the figure indicate exon positions with connecting thin dotted lines representing the borders of the intron. Numbers indicate branch point reads found in the combined RNA-seq datasets. Canonical branch points are shown in orange. C) Introns with alternative branch points: as in (B), but with alternative branch points shown in green.

For *trnK* and *trnA*, these alternative lariats utilize 5’ splice sites downstream of the canonical site, well inside the intron, suggesting a failure in 5’-splice site selection. The branch point A, however, is preserved in the alternative lariats of *trnK* for both investigated plant species and for *trnA* in tobacco, with only the 5’-splice site changing. This contrasts with *trnA* in *Arabidopsis*, where we identified an alternative lariat formed through an alternative branch point within exon 2. In the *atpF* intron, the branch point also shifted into exon 2 and the 5’-splice site moved into exon 1, resulting in a longer excised intron.

As an alternative method to verify the existence of alternative lariats and test for their abundance, we employed an RNase R assay. RNase R, a 3’-to-5’ exonuclease, can degrade non-branched precursors but is incapable of degrading lariats across the branch point (31). We analyzed RNase R treated total RNA through RNA gel blot hybridization using a probe for the *trnK* intron. This intron has the highest number of reads representing alternative lariats compared to canonical ones (Figure 2B). We analyzed RNA from both wild-type (wt) tobacco plants and a line that exhibits increased *matK* expression. This line, named 5’+HA, possesses an *aadA* selection marker cassette upstream of the *trnK* gene, with a 5’-extension of the *matK* reading frame encoding a Hemagglutinin (HA)-tag (19). This cassette is powered by the robust 16S rRNA promoter, which leads to readthrough into the downstream *trnK* gene, as indicated by an additional band of about 5 kb absent in the wt (Figure 3). Additional evidence supporting overexpression will be presented subsequently.

**Figure 3:**
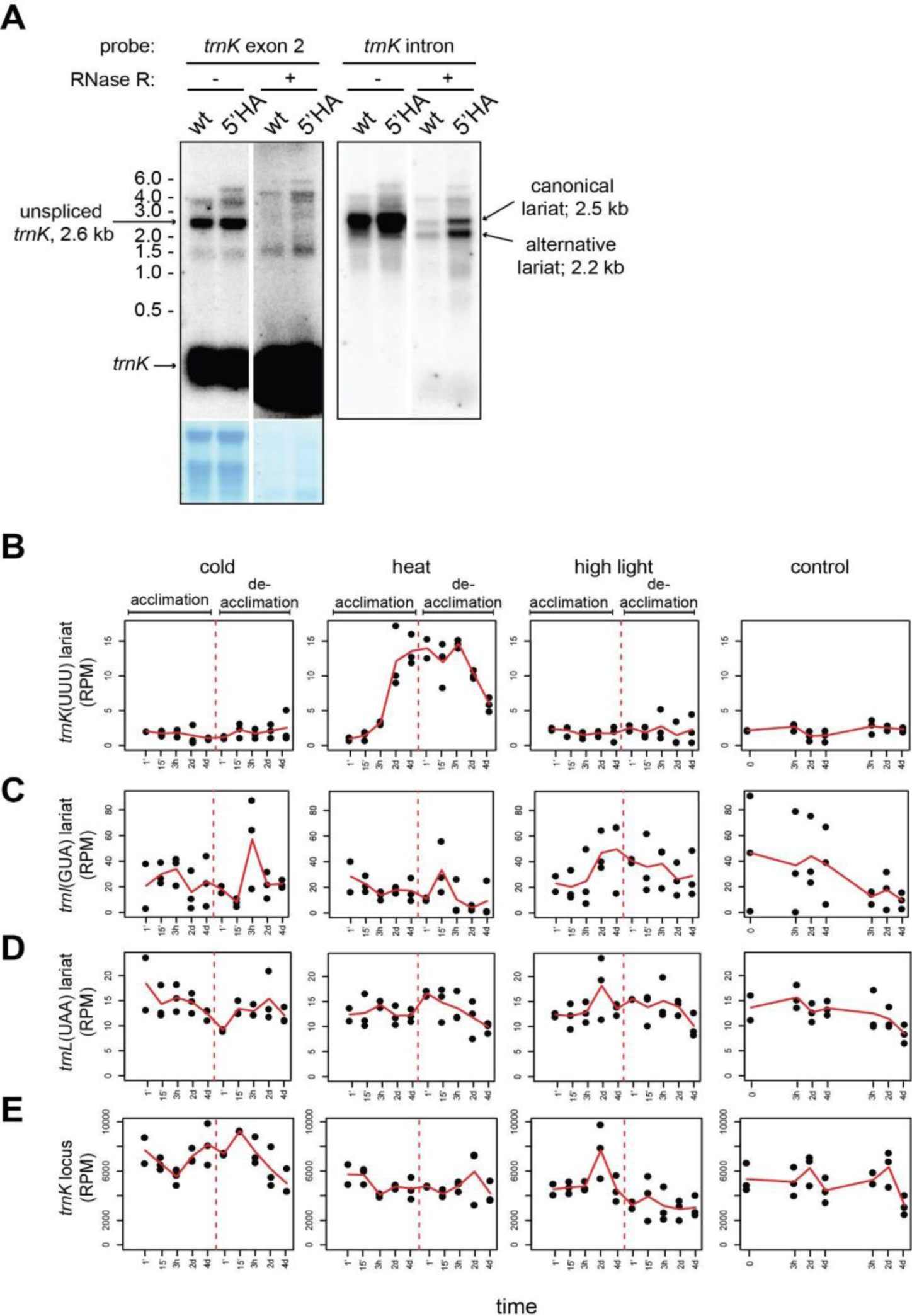
Accumulation of two *trnK* lariat isoforms. A) 5 µg of total plant RNA was either directly analyzed by RNA gel blot hybridization or after treatment with RNase R. Left: hybridization with a probe against exon 2 of *trnK*; methylene blue staining of the RNA gel blot to indicate equal loading. The white line indicates the removal of the marker lane and other lanes not relevant. Right: after stripping of the exon probe, a *trnK* intron probe was hybridized to the same membrane. B) Analysis of RNA-seq reads representing the aberrant *trnK* lariat isoform and canonical lariats in a dataset for cold, heat and high light acclimation (32). The dashed red line indicates the last sample taken in the initial acclimation phase before plants were put back to standard growth conditions for de-acclimation. C) Same as in (B), but for *trnI*(GUA) D) Same as in (B), but for *trnL*(UAA) E) Analysis of all RNA-Seq reads representing the *trnK* precursor RNA from the same dataset as used in B-C. Note that the increase in lariat reads during heat acclimation (B) is not reflected in an increase of precursor RNA.

Unspliced RNAs were detected using an exon 2 probe, revealing a major band of approximately 2.6 kb (Figure 3A). This signal for the linear *trnK* precursor RNA is entirely removed by RNase R treatment. The mature tRNA is enriched, as tRNAs are not an ideal substrate for RNase R (31), while rRNAs are also degraded efficiently (see Methylene blue stain below autoradiograms in Figure 3A). Conversely, using an intron-specific probe, we identified two transcripts that resist RNase R treatment, matching in size to the canonical and alternative lariats. Interestingly, the smaller, alternative lariat exhibits a stronger signal than the longer, canonical lariat (Figure 3A). This aligns with the RNA-seq experiment results, where more reads represent the alternative than the canonical lariat isoform. Together, the RNA-seq and RNA gel blot experiments confirm the existence of two alternative splicing products originating from the *trnK* precursor RNA.

Alternative splicing in the nucleus is a highly regulated process that responds to various external clues. To test whether alternative lariat formation is also reacting to environmental changes, we checked the accumulation of the lariat in recent RNA-seq data sets for *Arabidopsis thaliana*. We were using data from a study that exposed plants to mild changes in abiotic conditions requiring acclimation, but not yet exhibiting stress. The study used exposure to high light at 450 μmol photons m^−2^ s^−1^, cold exposure to 4°C, and heat exposure to 32°, all independently for four days, with subsequent de-acclimation for five days (32). In these datasets, we counted branch point reads for the alternative, aberrant *trnK* lariat (Figure 3B; reads for the canonical lariat are very rare) and also counted lariat reads for two further intron-containing tRNAs (Figure 3C, D). Neither in cold nor in high light did we find any significant changes in the accumulation of these lariats (Figure 3B-D). In heat acclimation however, a more than 7-fold increase in the alternative lariat was observed after three days of high temperature exposure that persisted for a few hours after return to regular growth temperature before it declined at the latest time points of de-acclimation (Figure 3B). The two other introns did not show such an increase in lariat reads. Also, the *matK* locus itself does not show increased expression during heat acclimation (Figure 3E). This indicates that heat is a specific modifier of *trnK* splicing that favors the formation of the aberrant lariat.

### The lysine codon read by tRNA-K differentiates between ribosomal and photosynthetic genes

To discern whether there is a bias in the distribution of the AAA codon read by tRNA-K(UUU) in different chloroplast genes, we conducted a hierarchical cluster analysis of all chloroplast protein-coding genes from tobacco based on their codon usage (Figure 4A). Two main clusters were found depending on their codon composition, which corresponded mainly to photosynthetic genes versus ribosomal genes. Fig 4B indicates that the lysine codon, AAA, is the main driver of this clustering, as it showed a significant (p value < 1.7*10^-9^, t-test) higher frequency in genes that code for ribosomal proteins, than encoding photosynthetic proteins.

**Figure 4:**
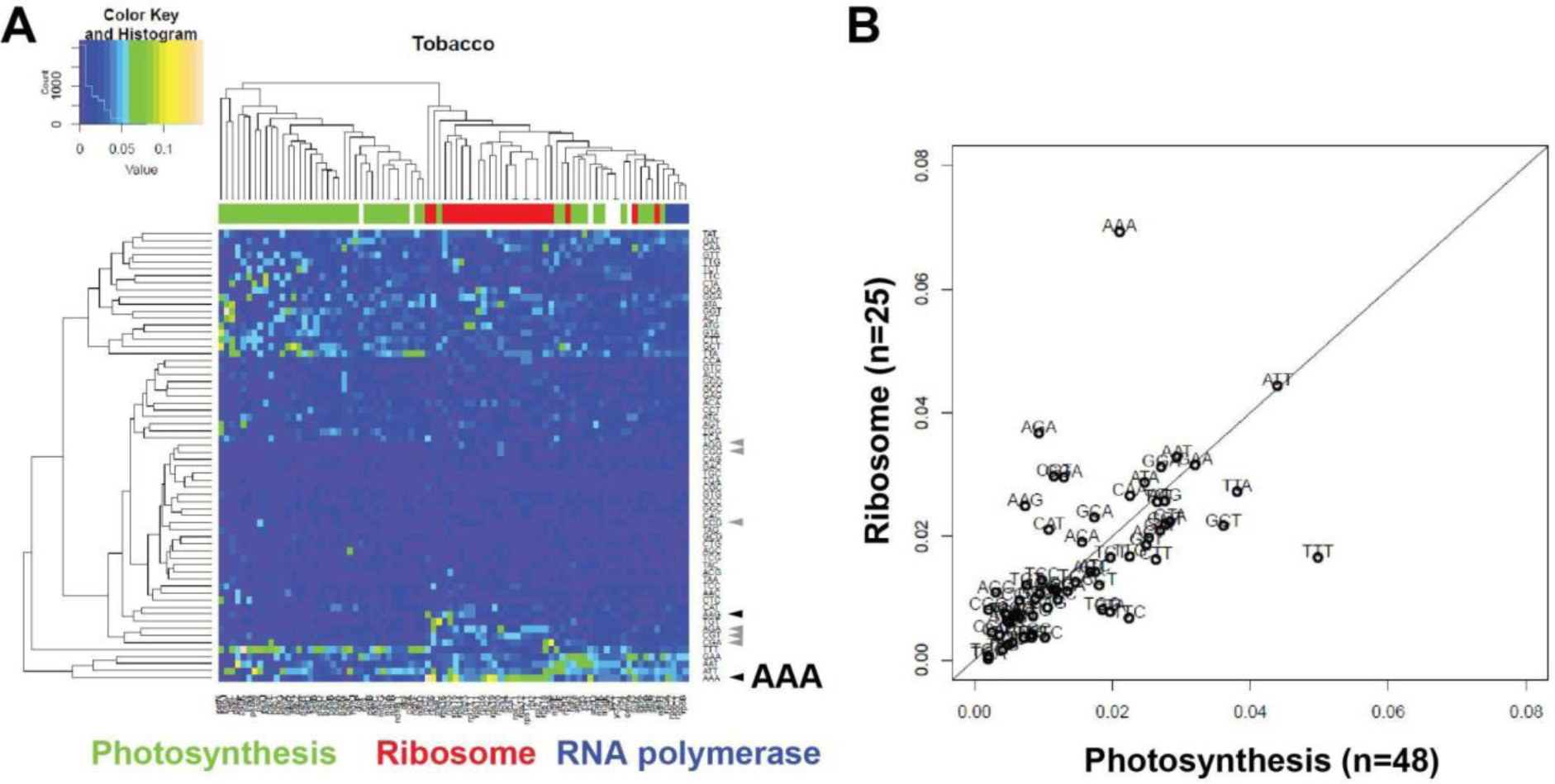
The AAA lysine codon is overrepresented in ribosomal protein genes. A) Hierarchical cluster analysis and heat map of the codon usage of each chloroplast gene. Chloroplast genes are functionally classified into three groups, photosynthesis, ribosome, and RNA polymerase. Selected codons relevant here are marked: black arrowheads: lysine codons; grey arrowheads: arginine codons. B) Comparison of codon frequencies in ribosomal versus photosynthesis genes.

Building on this discovery, we directly compared the codon frequencies in ribosomal protein genes versus photosynthesis genes. The AAA codon emerged as the most biased codon, with a marked preference for ribosomal protein genes. Given their function in binding rRNA, ribosomal proteins are known to be rich in positively charged amino acids, which generally necessitates a high presence of lysine codons. Although the same could be expected for arginine codons, we observed no such bias for arginine codons AGA, AGG, CGC and CGG, and only a mild overrepresentation of CGT and CGA codons in ribosomal protein genes. Considering the AT-rich nature of chloroplast genomes, a predilection for AAA codons could be partly explained by selection for AT-rich codons. However, the AAA codon certainly distinguishes between ribosomal and photosynthetic genes, leading us to hypothesize that this gene-group-specific bias might have regulatory implications. Intriguingly, the TTT codon, which codes for phenylalanine, displayed an opposite bias, being overrepresented in photosynthetic genes. Next, to investigate the potential importance of the AAA codon for the differential translation of gene groups, we sought methods to manipulate the levels of tRNA-K(UUU).

### Enhanced Levels of Mature tRNA-K in 5’+HA Transplastomic Plants

Among the two alternative splicing pathways for *trnK* that we identified, only one results in the production of mature tRNA-K. The alternative lariat pathway would fuse a significantly longer first exon to a standard second exon, which would lead to an aberrant, nonfunctional tRNA. However, we couldn’t detect this elongated tRNA in our RNA gel blots (Figure 3, exon probe hybridization), suggesting that the second splicing step may either be unsuccessful or the product is quickly degraded. Incorrect lariat formation might still play a regulatory role by competing with the generation of mature, correctly spliced tRNA-K.

We next asked whether there is a correlation between the expression levels of the *trnK* precursor / *matK* mRNA and the production of functional tRNA-K. We decided to explore this by using the previously generated transplastomic lines (19) that have *aadA* cassettes upstream or downstream of the *matK* reading frame, either with or without a triple HA-tag added to the coding sequence (Figure 5A). We used RNA gel blot hybridization to analyze the expression of *matK* in these lines. The 5’+HA line exhibits a significantly increased accumulation of the *trnK*-precursor relative to the wild-type (Figure 5C). Thus, the 5’+HA transplastomic line over-expresses the *trnK* precursor, essentially serving as an over-expressor of the *matK* mRNA.

**Figure 5:**
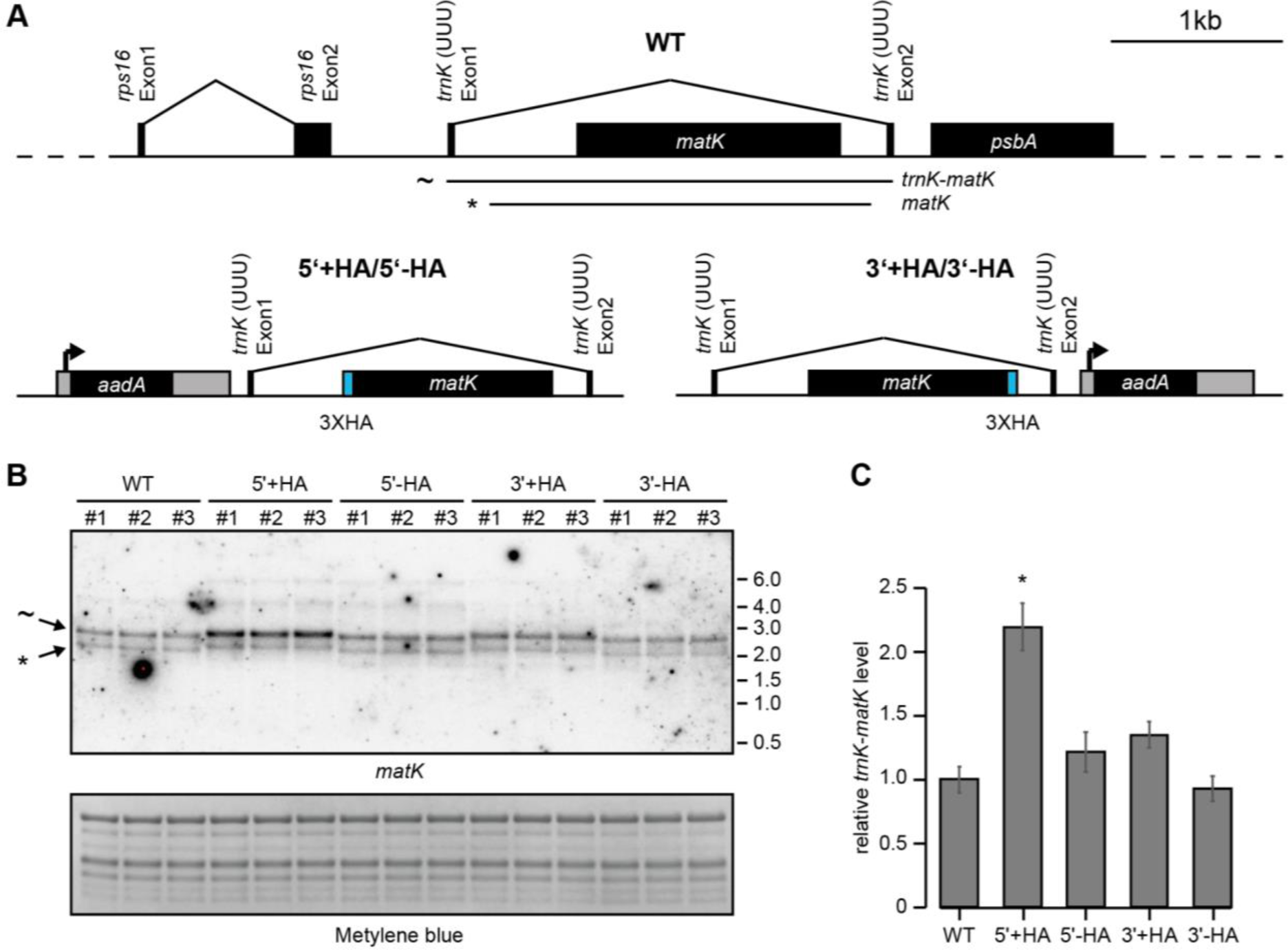
Analysis of unspliced *trnK* precursor transcript levels in transplastomic lines. A) Map of the *matK* genomic region in tobacco and a description of the alterations in the four transplastomic lines under investigation. Black boxes correspond to exons, and lines connecting exons correspond to introns. Gray boxes indicate positions of regulatory elements i.e. the tobacco 16S rDNA promoter and 3‘UTR of the Chlamydomonas *rbcL* gene. Blue boxes upstream (5‘+) or downstream (3‘+) of the *matK* open reading frame represent three copies of the hemagglutinin affinity tag (3XHA). 5‘- and 3‘-lines do not encode for this tag but still contain the *aadA*-cassette (spectinomycin-resistance gene) driven by the strong 16S rDNA promoter. Two different transcripts detected by RNA gel blot analysis representing the unspliced precursor as well as the lariat are indicated (∼ and *). B) RNA gel blot analysis of 3 µg of total tobacco plant RNA using a probe directed against the *matK* open reading frame. A methylene blue staining of the membrane before hybridization controls for equal loading. A band corresponding in size to the unspliced *trnK-matK* transcript (∼) and a band corresponding to the free *trnK* intron (*) are indicated. We interpret the latter as the non-canonical lariat. Note that the matK versions with 5’- and 3’-extended reading frames migrate slower in the gel than the wild-type and -HA lines. While 5’+HA lines shoes increased accumulation of RNAs, this was not observed for the 5’-HA line, suggesting that the HA-tag itself contributes to the overaccumulation of the precursor, potentially by altering RNA degradation rates. C) Quantification of the unspliced *trnK* transcript from the RNA gel blot analysis in B. Signal intensities were normalized against the mean of the three wild-type signals. Shown is the mean ± SD. The significance of differences was calculated using an ANOVA with Tukey HSD posthoc test. 5‘+ HA lines accumulate a significantly different amount of unspliced *trnK* compared to all other lines (p<0.05).

Next, we investigated whether the overexpression of the precursor also leads to increased levels of the mature tRNA-K. After separating small RNAs from the transplastomic lines via PAGE, we hybridized the blots with probes against tRNA-K and two other tRNAs, tRNA-L-UAA and tRNA-V-UAC (Figure 6A). *trnL* contains a group I intron that does not depend on MatK, while *trnV* contains a group IIA intron, but was not significantly enriched in our RIP-Seq experiment (Figure S1). We observed a significant increase in tRNA-K levels relative to wild-type in 5’+HA plants, while the abundance of the two other tRNAs remained unchanged across the analyzed genotypes (Figure 6B).

**Figure 6:**
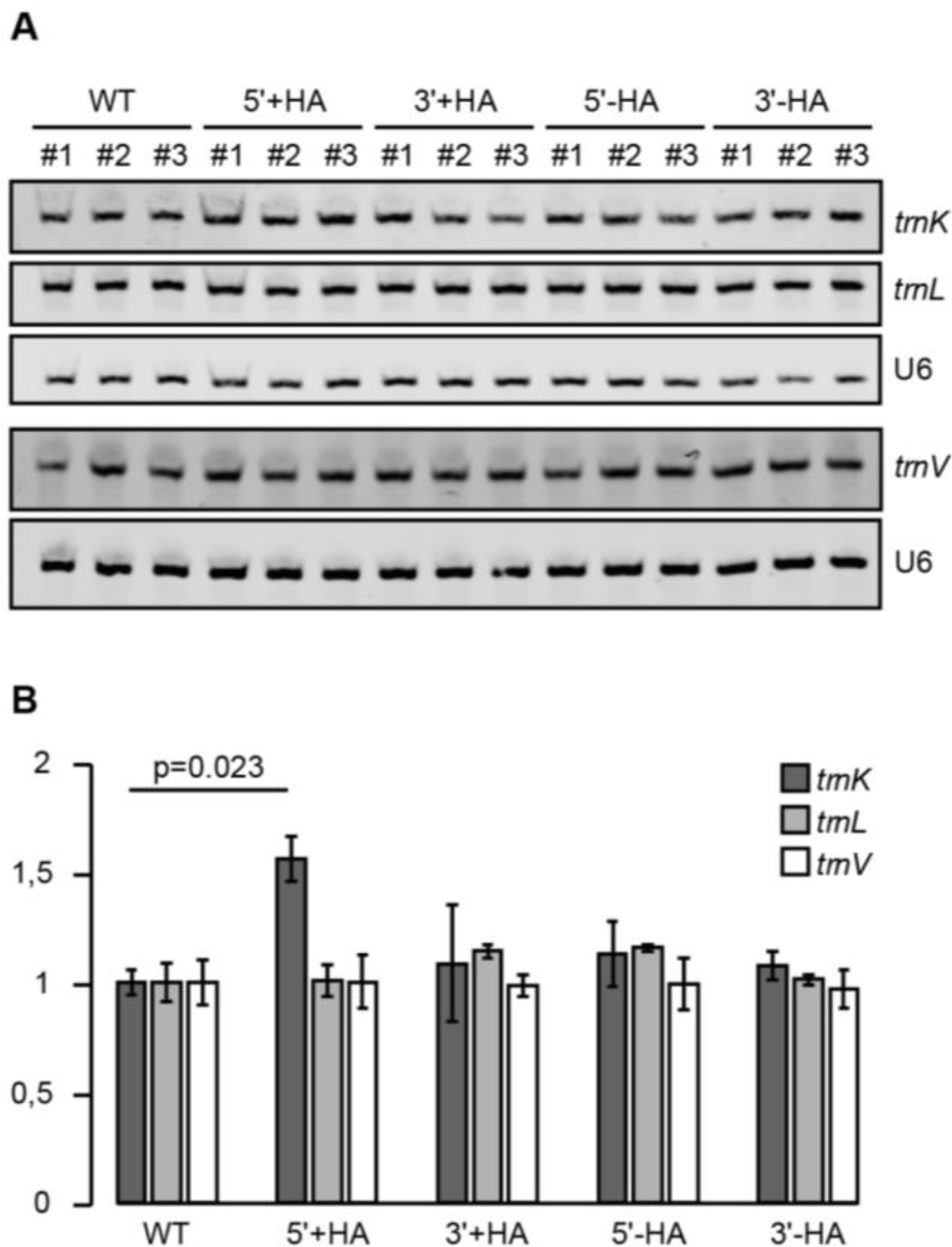
*trnK* over accumulates in the 5‘+HA line. A) RNA gel blot analysis of selected intron-containing chloroplast tRNAs. 2µg total RNA was separated on denaturing polyacrylamide gels and transferred to a nylon membrane. Chloroplast tRNAs were probed with fluorescently labeled oligonucleotides complementary to the second exon. A probe detecting U6 snRNA was used as a loading control. This experiment was conducted with three biological replicates (plants #1-3). B) Quantification of signal intensities from RNA gel blot analysis in B. The significance of differences was calculated using an ANOVA with Tukey HSD posthoc test (significance threshold: p<0.05).

### Increased *matK*/tRNA-K(UUU) levels lead to shifts in the translation of PG versus GA genes

To investigate the impact of increased expression levels of tRNA-K(UUU) in 5’+HA lines on translation, we conducted a ribosome profiling experiment. We compared 5’+HA plants with 5’-HA plants by subjecting leaf tissue from 14-day-old tobacco plants to ribosome footprint purification, followed by fluorescent dye labeling of the footprints. These labeled footprints were then hybridized to a high-resolution tiling microarray, which covers all chloroplast ORFs, as previously described (33). This process allowed us to map ribosome footprints with a resolution of approximately 30 nucleotides, thanks to the 20-nucleotide overlap of the 50-mers on the array. We summed up the array signals of all probes for chloroplast genes encoding photosynthetic proteins as well as genes encoding ribosomal proteins and compared the distribution of translation scores. We observed a significant up-shift in the ribosomal genes compared to the photosynthetic genes, indicating an overall increase in ribosome occupancy on ribosomal protein mRNAs in the *matK/trnK* overexpressor line (Figure 7A).

**Figure 7:**
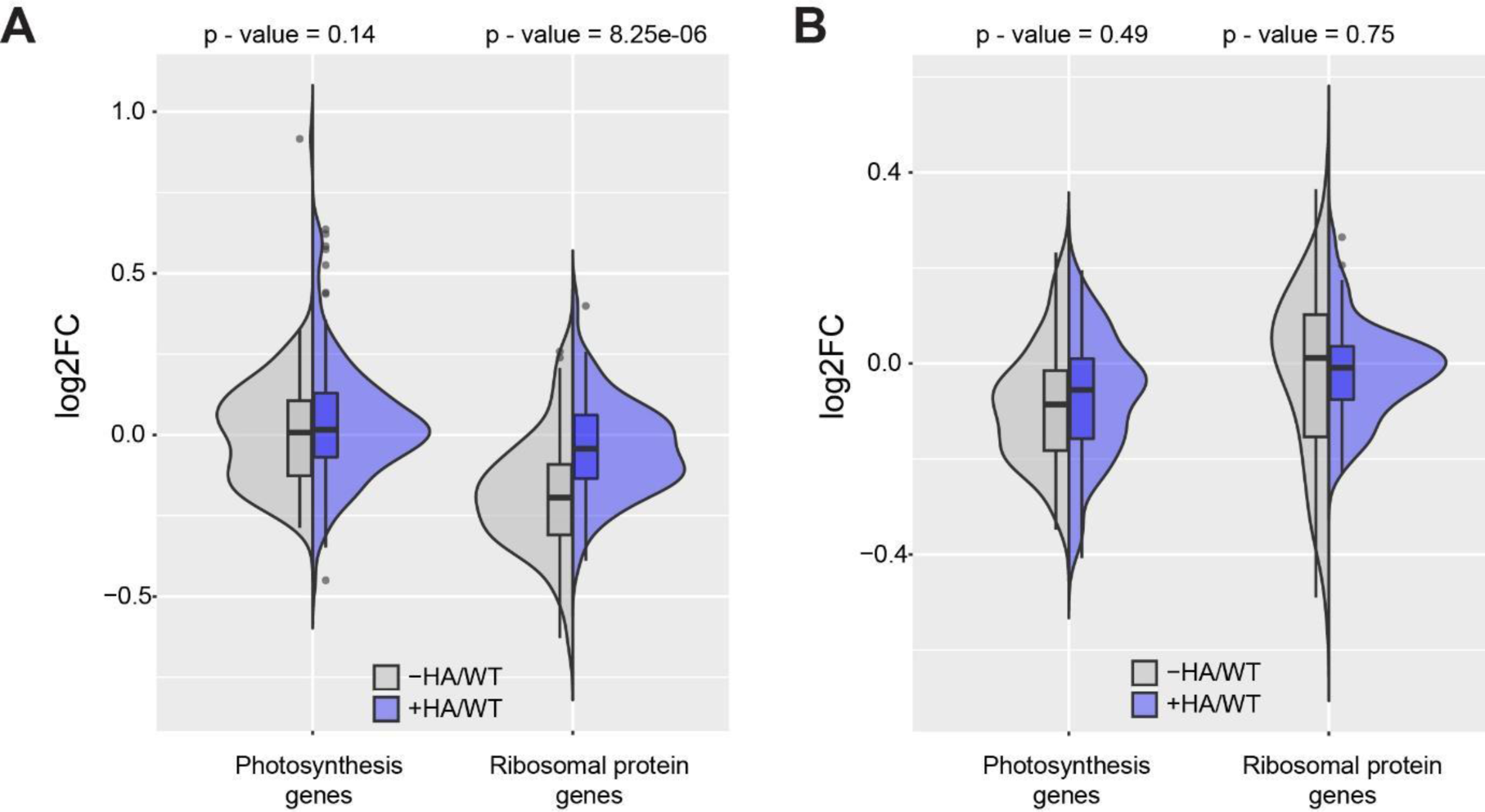
MatK levels correlate with translational changes of GA versus PS genes. A) Comparison of chloroplast RNA accumulation between *matK/trnK*-overexpressing plants (+HA) and control plants (-HA) in comparison to wild type (wt) using microarray-based RNA profiling. The microarray data were separately analyzed for genes encoding photosynthetic proteins and genes responsible for ribosomal proteins. The p-values provided indicate the significance of the difference in RNA accumulation between the overexpressor and control plants for each gene group. Importantly, our analysis revealed a significant difference in ribosome footprints between genes encoding photosynthetic proteins and genes encoding ribosomal proteins when comparing the 5’+HA *matK/trnK* overexpressor lines with the control lines. B) Same analysis as in (A) but for RNA from the same plant lines. Note that genes for ribosomal proteins show no significant increase in RNA levels in the overexpressor for neither photosynthetic nor ribosomal protein genes.

To determine whether this shift in ribosome occupancy is caused indirectly by an increase in the respective mRNA levels, we analyzed RNA accumulation by a parallel microarray hybridization experiment. Here, neither the photosynthetic genes nor the genes coding for ribosomal proteins show a change in signal in the overexpressor versus controls (Figure 7B). In conclusion, increasing the expression of *matK/trnK* correlates with an increase in translation of ribosomal protein mRNAs.

## Discussion

Translation has emerged as a key point for chloroplast gene regulation (6). It has been found that chloroplast translation rates provide a better representation of the subunit stoichiometry in thylakoid membrane complexes involved in photosynthesis compared to RNA levels (34, 35). Furthermore, chloroplast translation efficiency responds rapidly to changes in light, temperature and developmental stage (36–40). However, the mechanisms underlying translation adjustments are still poorly understood.

So far, very few general mechanisms of translational regulation have been observed in chloroplasts, and all of them are based on similarities to bacterial systems, like ppGpp based signaling, the role of PSRP1 and redox-regulated translation elongation factor Tu (41–45). In addition to these general regulatory mechanisms of translation, there are also factors that are specifically supporting translation of to one or few mRNAs (Prikryl et al., 2011; Hammani et al., 2012; summarized in Zoschke and Bock, 2018), but these do not account for the co-regulation of gene groups at the translational level. Here, we propose a simpler explanation for co-translation of mRNAs, which involves modulating the abundance of a specific tRNA species that recognizes codons overrepresented in a particular gene group.

We have discovered that the AAA codon is the most biased codon between photosynthesis and ribosomal genes. Increasing the levels of tRNA-K(UUU) in an over-expressor of the *trnK/matK* locus correlated with an elevation of ribosomal protein gene translation, as observed through ribosome profiling. We note however, that other functions of MatK could contribute to driving the translation of ribosomal proteins. MatK is also required for splicing of ribosomal protein mRNAs, such as *rps12* and *rpl2*. It is conceivable that the production of ribosomal proteins via MatK-dependent splicing contributes to an increased translational capacity in the *trnK/matK* over-expressor. To determine the contribution of splicing efficiency of different *matK* targets to the translation enhancement of mRNAs for ribosomal proteins, further investigation is needed, such as ectopic expression of a pre-spliced version of *trnK* to separate its expression from the influence of MatK. At present however, we propose the most simple and parsimonious model that MatK impacts activation of translation of ribosomal mRNAs by supporting canonical lariat formation and thus tRNA-K(UUU) production (Figure 8).

**Figure 8:**
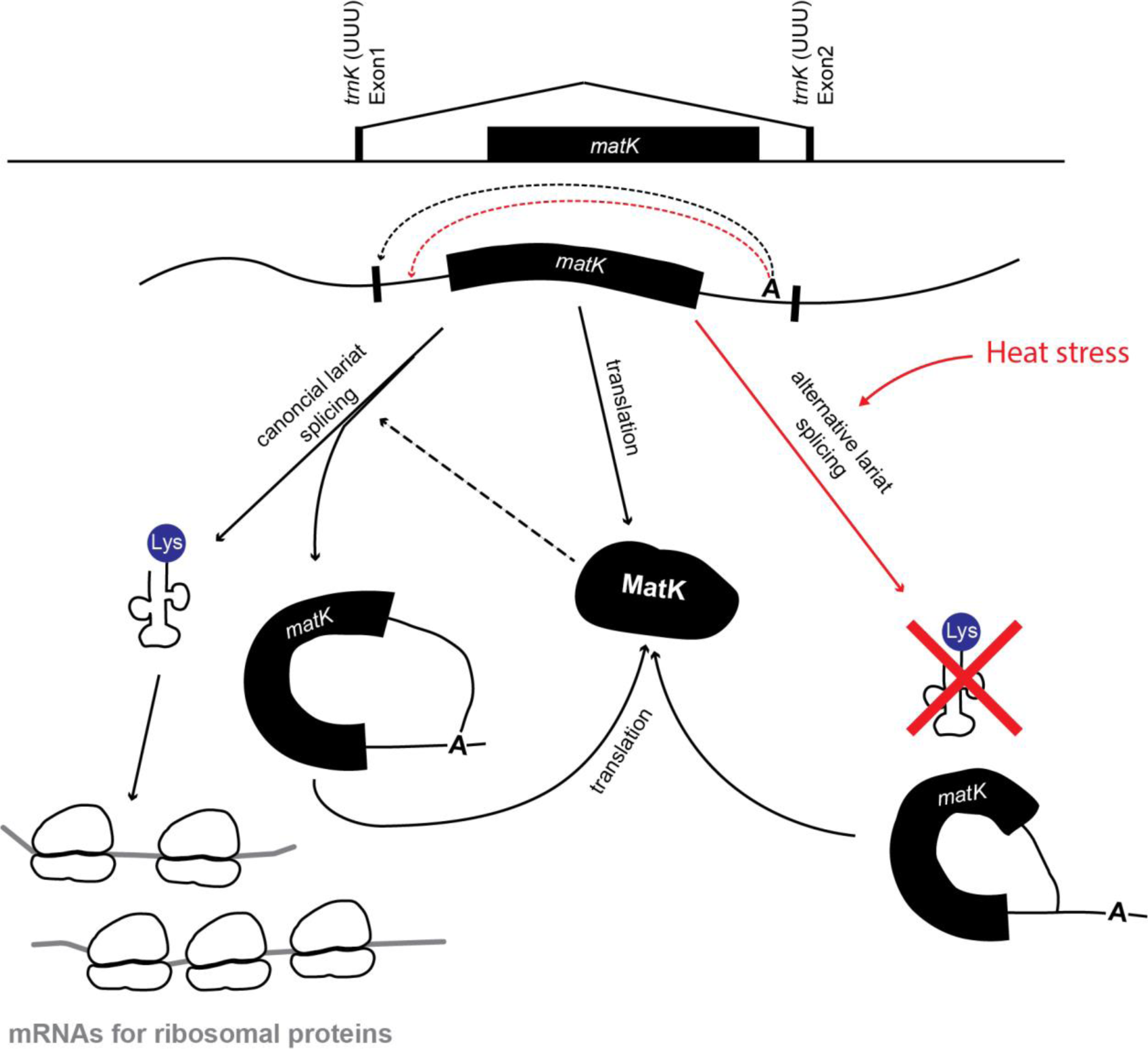
Model of the impact of alternative lariat formation on gene-group regulation. There are two possible lariat isoforms in the trnK intron, both originating from the same branch point A, but with different 5’-splice site. The canonical lariat (black dotted line originating from branch point A) leads to functional tRNA-K production and is supported by MatK. The tRNA is serving AAA codons that are particularly frequent in mRNAs for ribosomal proteins, thus leading to increased translation of ribosomal proteins versus photosynthetic proteins. Alternative lariat formation (red dotted line originating from branch point A) is induced by heat acclimation and does not lead to functional tRNA.

This model is reminiscent of the translation of stress-related genes in yeast that tend to use rare codons (46) – under stress, tRNA pools are skewed toward those tRNAs that recognized rare codons so that stress proteins are translated more efficiently than protein involved in growth and proliferation (47). In general, differential tRNA expression is a well-known phenomenon (48, 49), but was rarely connected to functional readouts, e.g. in cancer progression (50). This is probably due to the high redundancy of isodecoders in the nucleus, which in most cases prevents a simple genetic approach for functional studies. For example, a knock-out of six tRNA-Phe genes was required to uncover the importance of this isodecoder set for mouse embryo development (51), but at the same time revealed that one particular tRNA-Phe gene was relevant alone for neuronal function (51). Such single-isodecoder effects are however rare. The chloroplast offers the interesting situation of a small, minimal set of tRNAs with little isoacceptor redundancy. Any manipulation of single tRNAs can directly affect translation and compensation by other tRNAs is minimal (52–55). The manipulation of tRNA-K by MatK overexpression demonstrates that specific translation changes can be realized by single tRNA species. Whether more tRNA species entail a clear effect on just a subset of chloroplast mRNAs or result in global translational changes, remains to be determined.

Regulating translation through tRNA-K(UUU) abundance can also resolve the longstanding issue of how genes in mixed operons can be differentially regulated. Unlike bacterial operons, chloroplast operons are typically mixed, meaning that ribosomal protein genes are co-transcribed with photosynthesis genes, as seen in the *psaA/psaB/rps14* operon. Initially, it was believed that the processing of such mixed precursor RNAs into monocistronic forms would determine the eventual translation of an open reading frame. However, polysome analyses and ribosome profiling experiments have demonstrated that the processing state of an mRNA has little impact on the translation rate in most cases (1, 34, 35, 56, 57). On the other hand, tRNA-K(UUU) abundance is completely independent of processing state and adjacent ORFs, enabling regulation of ribosomal protein genes across highly diverse contexts and transcripts in a straightforward manner. This could be for instance relevant during leaf development. In meristematic and young leaf tissue, the gene expression apparatus is primarily expressed to prepare for the later production of the photosynthetic machinery (1). We propose that an early expression of MatK and thus tRNA-K(UUU) during leaf development will contribute to setting up the gene expression apparatus, in particular the ribosome. This temporal separation of the expression of two gene groups is logical as chloroplast translation is particularly crucial during chloroplast biogenesis, i.e. in young tissue. By contrast, once established, the photosynthetic apparatus remains, with very few exceptions, highly stable (58, 59) and the demand for translational capacity is accordingly low later in development (1).

Furthermore, it is important to note that plastid translation remains relevant even in the absence of photosynthesis due to the presence of reading frames essential for non-photosynthetic metabolism. One example is *accD*, which encodes an essential enzyme involved in the initial step of fatty acid biosynthesis. Consequently, we predict that in roots and other non-photosynthetic tissues, the expression of *trnK/matK* will contribute to maintaining basal levels of the translational apparatus, ensuring the proper synthesis of proteins necessary for non-photosynthetic metabolic processes. In addition to potential developmental relevance, we present evidence that *trnK* splicing is changed during heat acclimation with more aberrant lariats being formed at high temperatures. How this impacts tRNA abundance and translational output during heat response remains to be determined.

### MatK as a modifier of alternative lariat formation of the tRNA-K (UUU) precursor RNA has a potential relevance during temperature acclimation

Three routes have been described for group II intron splicing - the canonical branching pathway, the hydrolytic pathway without lariat formation and the circularization pathway (60). None of these routes is compatible with the alternative lariats identified here. In the absence of functional evidence for the alternative splicing products, we consider the alternative lariat as aberrant and detrimental, possibly arising due to misfolding of the RNA that is enhanced at higher temperatures. This would be reminiscent of the finding that the circle-to-lariat ratio of the Ll.LtrB group II intron from *Lactococcus lactis* is significantly influenced by temperature changes (61) and the general sensitivity of splicing to heat and cold in plants (62). MatK is fostering the canonical splicing pathways as suggested by the correlation of its expression with tRNA-K(UUU) levels in the over-expressor. Our RIP-Seq results furthermore point out that the primary targets within introns are DI and DVI. This is in line with bacterial maturases that are known to deeply penetrate into DI, which supports the IBS-EBS interaction (63) and that exert their influence on splicing by engaging in weak interactions with the intron’s active site and DVI (23, 64). These studies strongly suggest that bacterial maturases play a crucial role in modulating splicing dynamics by directly interacting with key regions within the intron, such as the active site and the branch site adenosine residing in DVI. We hypothesize that MatK suppresses the formation of an alternative structure favored at higher temperature that allows the selection of a faulty 5‘-splice site. This can be tested by structure probing experiments of the *trnK* intron in the presence and absence of MatK. Our findings indicate that MatK regulates chloroplast gene expression beyond functioning as a general splicing factor, and that control of abundance of specific tRNAs adds an exciting layer to the regulatory hierarchy controlling translation in chloroplasts.

## Material and Methods

### Plant material

Nicotiana *tabacum* transplastomic lines expressing Hemagglutinin (HA)-tagged MatK as well as control lines were prepared previously (19). Plants were grown on soil under long day conditions (16 hours light, 8 hours dark) at 27°C with light intensities of approximately 300 μmol. m-2. s −1. For acquisition of 7-day-old tissue for chloroplast isolation, seeds were sown on polyamide nets (mesh size 500 μM, Franz Eckert GmbH) on soil. Both seed sowing and seedling harvesting were performed at 10:00 am.

### Chloroplast isolation and stroma extraction

The procedure of chloroplast isolation from tobacco was modified from a protocol developed for maize (65). Seedlings of 7-day-old tobacco (without roots) were harvested from 3-4 plates grown on mesh-covered soil (d = 14 cm) and were homogenized with a waring blender in 350 ml grinding buffer (50 mM HEPES-KOH (pH 8.0), 330 mM Sorbitol, 2 mM EDTA, 1 mM MgCl2, 1 mM MnCl2, 0.25% BSA, 1.5 mM Sodium ascorbate), once for 5 s at low speed, and twice more for 5 s at the high-speed setting. Next, the mixture was filtered through one layer of MicroCloth (Calbiochem) and centrifuged at 1000 g for 6 min, after which the pellet was resuspended in 1-2 ml of HS buffer. After adding 3-4 volumes of HS buffer, it was centrifuged at 1000 g for 6 min. The supernatant was discarded again, 200-400 μl EX buffer (0.2 M KAc, 30 mM HEPES-KOH (pH 8.0), 10 mM MgAc2, 2 mM DTT, before use, add 1× protease inhibitor (Protease Inhibitor Cocktail Tablets, Roche), 0.4 mM PMSF, 0.1 μg/ml Aprotinin) was added to the pellet, and the mixture was squeezed through a syringe (needle: 0.55 × 25 mm) ∼40 times in order to lyse chloroplasts. Finally, lysed chloroplasts were centrifuged at 30,000 g for 30 min to separate the membrane and stroma fractions. The stroma fraction was used for immunoprecipitation (IP) of MatK.

### RIP-Seq

For each MatK co-IP, two volumes of co-IP buffer (Co-IP buffer: 0.15 M NaCl, 20 M Tris-HCl (pH 7.5), 2 mM MgCl2, 0.5% NP-40 (v/v)) and 5 μg of mouse anti–HA antibody (Sigma) were added to the stroma fraction containing approximately 200 μg of protein. The stroma-antibody mixture was rotated at 12 rpm for 1 hour, after which 50 μl Magnetic Beads (Life Technologies) were added and the reaction was rotated for one more hour. Next, the beads were pelleted on a magnetic rack, and the supernatant was taken for RNA extraction and Western blot. Finally, the IP pellet was washed 3 times with co-IP buffer, and 200 μl EX buffer was added to the IP pellet before storage or RNA extraction. The same fraction (1/50) was aliquoted from both IP supernatant and pellet for Western blot.

RNA was extracted with Phenol/Chloroform from the IP pellet of HA tagged and control lines. Next, the RNA samples were used for library construction using ScriptMiner Small RNA-seq library preparation Kit (Epicentre) and sequencing was performed on an Illumina NextSeq500.

### Mapping

Quality-trimmed Illumina reads from the RIP-seq experiment were mapped to the *Nicotiana tabacum* chloroplast genome (NC_001879.2) with the repeat region masked. We utilized CLC Workbench (Version 6.0.1) with the following settings: Mismatch cost: 2; Insertion cost: 3; Deletion cost: 3; Length fraction: 0.5; Similarity fraction: 0.8. Local alignment was applied to enhance accuracy.

### RIP-Seq analysis - Domain-Based Peak Selection

To identify the preference of MatK for domains of type-2-introns (66), we sorted the bases within each domain based on their normalized +HA to -HA ratios and selected the top 40 bases. The mean of the selected bases was used for heatmap generation, utilizing the ComplexHeatmap library (67). Domains were clustered using the Euclidean distance method.

### RNA gel blot hybridization

Total RNA was extracted using TRIzol (Invitrogen) following the manufacturer’s protocol. Detection of RNAs with radiolabeled probes was performed as described (68). For detecting RNAs encompassing *trnK* exon sequences, the trnKex2 oligonucleotide (tgggttgcccgggattcgaacccggaactagtcgg, Figure 3A) was directly end-labeled using T4 polynucleotide kinase. For subsequent detection of all intron-containing transcripts, including free introns after splicing, oligonucleotides MatKseqfor1 (gaagttcgatacccttgttcc) and matKolrev1 (gataattcgccagatcattgataca) were used for amplifying part of the *matK* reading frame. The resulting PCR product was body-labeled, denatured and hybridized to the stripped RNA gel blot (Figure 3A).

Mature tRNAs were separated by denaturing Urea-PAGE, transferred to nylon membranes and hybridized in ULTRAhyb™-Oligo buffer (Invitrogen). DNA oligonucleotides used as probes were extended with 5-Azido-PEG4-dCTP (Jena Bioscience) and terminal transferase (NEB). Purified extended oligonucleotides were click-labelled with Cyanine5.5 or Cyanine7.5-alkyne (Lumiprobe). Probe sequences for Figure 6A were designed for *trnK* exon 2 (trnKex2: tgggttgcccgggattcgaacccggaactagtcgg); *trnV* exon 2 (trnVex2: tagggctatacggactcgaaccgtagaccttctcg); *trnL* exon 2 (trnLex2: tggggatagagggacttgaaccctcacgatttttaaagtcgacggatttt); and U6 (aggggccatgctaatcttctctgtatcgttccaa). For probe generation against *matK* transcripts (Figure 5B). Oligonukleotides matKNt.rp (cattcccgtgactttatgggc) and matKNt.T7 (taatacgactcactatagggaagaagctcgtgggaaggtc) were used to generate PCR fragments that served as templates for *in vitro* transcription reactions in the presence of radio-labeled nucleotides (Figure 5B).

### Immunoblot

Immunoblot analyses were performed as described (27).

### Ribosome profiling

Ribosome profiling started from seven day old material and was done as described based on microarray hybridization of an oligonucleotide array covering the entire tobacco chloroplast coding sequence with 50mer oligonucleotides on both strands (33). The data can be found in supplemental dataset 1.

## Acknowledgments

We thank Ines Gerlach for excellent technical support. This research was supported by the DFG via TR175-A02 to CSL, TR175-C05 to K.K., TR175-D01 to U.O., TR175-A07 to H.R. and TR175-A04 and ZO 302/5-1 to R.Z.

## Data availability

Sequencing data have been deposited in GEO, accession number GSE245847.

